# Leveraging brain cortex-derived molecular data to elucidate epigenetic and transcriptomic drivers of neurological function and disease

**DOI:** 10.1101/429134

**Authors:** Charlie Hatcher, Caroline L. Relton, Tom R. Gaunt, Tom G. Richardson

## Abstract

Integrative approaches which harness large-scale molecular datasets can help develop mechanistic insight into findings from genome-wide association studies (GWAS). We have performed extensive analyses to uncover transcriptional and epigenetic processes which may play a role in neurological trait variation.

This was undertaken by applying Bayesian multiple-trait colocalization systematically across the genome to identify genetic variants responsible for influencing intermediate molecular phenotypes as well as neurological traits. In this analysis we leveraged high dimensional quantitative trait loci data derived from prefrontal cortex tissue (concerning gene expression, DNA methylation and histone acetylation) and GWAS findings for 5 neurological traits (Neuroticism, Schizophrenia, Educational Attainment, Insomnia and Alzheimer’s disease).

There was evidence of colocalization for 118 associations suggesting that the same underlying genetic variant influenced both nearby gene expression as well as neurological trait variation. Of these, 73 associations provided evidence that the genetic variant also influenced proximal DNA methylation and/or histone acetylation. These findings support previous evidence at loci where epigenetic mechanisms may putatively mediate effects of genetic variants on traits, such as *KLC1* and schizophrenia. We also uncovered evidence implicating novel loci in neurological disease susceptibility, including genes expressed predominantly in brain tissue such as *MDGA1, KIRREL3* and *SLC12A5*.

An inverse relationship between DNA methylation and gene expression was observed more than can be accounted for by chance, supporting previous findings implicating DNA methylation as a transcriptional repressor. Our study should prove valuable in helping future studies prioritise candidate genes and epigenetic mechanisms for in-depth functional follow-up analyses.

## Background

Genome-wide association studies (GWAS) have been integral in identifying thousands of genetic variants associated with complex traits and disease. The vast majority of genetic variants identified in these studies reside in intergenic or intronic regions of the genome and are therefore predicted to exert their effects on complex traits via changes in gene regulation^1^. Furthermore, there is evidence which suggests that GWAS hits are often located within regions of open chromatin and enhancers^2^. Typically, genetic variants associated with molecular phenotypes are enriched amongst SNPs that are linked to traits and diseases^3^. Such variants are known as quantitative trait loci (QTL) and can affect molecular phenotypes such as: gene expression (eQTL), and epigenetic mechanisms including DNA methylation (mQTL) and histone acetylation (haQTL). DNA methylation and histone acetylation are alterations that affect gene expression without altering the DNA sequence. Several genetic variants have been identified that occur in the same genomic region and influence both gene expression and DNA methylation. In these cases, it is possible that the eQTL and mQTL share a common causal variant (CCV)^4^.

Several post-GWAS approaches exist to help functionally characterise non-coding variants^5–7^. In particular, there has been an emphasis on integrating eQTL and GWAS data together, which can be valuable in terms of identifying the underlying genes responsible for associations detected by GWAS. Recently, similar endeavours have extended the scope of their analysis to also evaluate additional molecular phenotypes (e.g. mQTL and haQTL) as well as gene expression^8–11^. A novel method in this paradigm involves calculating approximate Bayes factors^12^ to assess the likelihood that the genetic variants responsible for an association with a complex trait is also responsible for influencing intermediate molecular phenotypes (i.e. the likelihood they share a CCV). This multiple-trait colocalization (moloc) method has been shown to help characterise GWAS loci and develop mechanistic insight into the causal pathway from genetic variant to complex trait^13^. Furthermore, inclusion of an additional molecular trait into the analysis (e.g. complex trait, gene expression and DNA methylation vs. complex trait and gene expression alone) has been shown to increase power and assist in identifying novel disease susceptibility loci^13^.

The recent large influx of tissue specific molecular data provides an unprecedented opportunity to assess the functional relevance of GWAS hits. Recently, a resource has become available that comprises QTL data derived from the dorsolateral prefrontal cortex in up to 494 subjects^14^. Brain xQTL Serve provides a list of SNPs associated with gene expression, DNA methylation and/or histone modifications specific to the same brain region^14^. Whilst progress has been made in terms of identifying genetic variants influencing neurological phenotypes and diseases, not enough is known about the biological effects of genetic risk factors. In this study, we have jointly analysed genetic variants identified from GWAS of 5 neurological traits and diseases alongside the variants listed in the Brain xQTL Serve resource. In doing so, we aim to identify CCVs for neurological traits and gene expression, and where possible, DNA methylation and histone acetylation. Uncovering evidence that epigenetic factors reside on the causal pathway along with gene expression can be extremely valuable for disease prevention due to early diagnosis.

## Methods

### Neurological genome-wide association studies

We obtained summary statistics from 5 independent GWAS for the following neurological traits: Neuroticism (n=274,108)^15^, Schizophrenia (cases=35,467, controls=46,839)^16^, Educational Attainment (n=293,723)^17^, Alzheimer’s disease (cases=17,008, controls=37,154)^18^ and Insomnia (n=336,965)^15^. Information on all GWAS datasets can be found in Supplementary Table 1. Linkage disequilibrium (LD) clumping was undertaken using PLINK v1.9^19^ with a reference panel consisting of European (CEU) individuals from phase 3 (version 5) of the 1000 genomes project^20^. This allowed us to identify the top independent loci for each set of results based on the conventional GWAS threshold (P<5.0×10^−08^).

### Brain-tissue derived quantitative trait loci for three molecular phenotypes

All QTL data used in this study were obtained from the Brain xQTL Serve resource^14^. Genotype data in this resource was generated from 2,093 individuals of European descent from the ROS and MAP study cohorts (http://www.radc.rush.edu/). Gene expression (RNA-seq; n=494), DNA methylation (450K Illumina array; n=468) and histone modification (H3K9Ac ChIP-seq; n=433) data were derived from the dorsolateral prefrontal cortex of post-mortem samples. eQTL were based on 13,484 expressed genes, mQTL on 420,103 methylation sites and haQTL on 26,384 acetylation domains. eQTL and haQTL results were available for variants within 1MB of their corresponding probes, whereas mQTL results were restricted to a 5kb window^14^.

### Gene-centric multiple-trait colocalization

We extracted effect estimates for all variants within 1MB of the lead SNP for each clumped region using results from each of the 5 GWAS. P-values for molecular QTL were then extracted for the same set of SNPs using the Brain xQTL resource. Loci residing within the Major Histocompatibility Complex (MHC) region (chr:6 25Mb –35Mb) were removed due to extensive LD within this region which may result in false positive findings. The moloc method was then used to assess the likelihood that the variant at each region responsible for variation in complex traits was also responsible for influencing the expression of a nearby gene (i.e within a 1MB distance of the lead GWAS SNP). As demonstrated previously^13^, we simultaneously investigated whether variants responsible for both gene expression and complex trait variation may also influence proximal epigenetic traits in a gene-centric manner. However, unlike previous work which evaluated 3 traits at a time, we have investigated up to 4 traits in each analysis (i.e. complex trait, gene expression, DNA methylation and histone acetylation).

To achieve this, we used coordinates from Ensembl^21^ to map CpG sites and histone peaks to genes using a 50kb window upstream and downstream of each gene. We then ran moloc to assess all Gene-CpG-Histone combinations within each region of interest. Summed posterior probabilities were computed for all scenarios where GWAS trait and gene expression colocalized. The reason for this is because if epigenetic mechanisms are responsible for mediating the effect of genetic variants on complex traits then we would expect gene expression to also reside on this causal pathway.

Therefore, 10 scenarios were considered of interest; GE, GE,M, GE,H, GE,M,H, GEM, GEM,H, GEH, GEH,M, GEMH, where evidence of a shared causal variants for GWAS complex traits is defined as ‘G, gene expression as ‘E’, DNA methylation as ‘M’ and histone acetylation as ‘H’. The ‘,’ denotes a scenario where there is a different causal variant for each molecular phenotype. For example, GE,M would represent a situation where the same causal variant is shared between the GWAS trait and gene expression, but a different causal variant for DNA methylation.

As recommended by the authors of moloc^13^, a summed posterior probability of association (PPA) >= 80% for these 10 scenarios was considered strong evidence that a genetic variant was responsible for changes in both molecular phenotype(s) and neurological trait variation. Therefore, a GEMH scenario with a posterior probability >=80% would represent a case where there is evidence that GWAS trait, gene expression, DNA methylation and histone acetylation colocalize and share a causal variant. When a Gene-Trait combination provided evidence of colocalization with multiple CpG sites or histone peaks, we only reported the association for the combination with the highest PPA. This was to reduce the number of findings detected due to co-methylation/probes within the same histone peak that were measuring the same epigenetic signatures.

Regions with fewer than 50 common SNPs (MAF <=5%) were not considered in the moloc analysis in order to reduce the number of spurious findings. Prior probabilities of 1×10^−04^, 1×10^−06^, 1×10^−07^ and 1×10^−08^ were used in all analyses which was also recommended by the authors of moloc. Furthermore, we used the option to adjust Bayes factors for overlapping samples as this was the case for the xQTL datasets. Manhattan plots to illustrate findings were subsequently generated using code adapted from the ‘qqman’ package^22^.

### Identifying potentially novel loci in disease susceptibility

We also applied our analytical pipeline as described above to independent GWAS loci with p-values between the conventional threshold (P<5.0×10^−08^) and P≤1.0×10^−06^. All parameters were the same as in the previous analysis. We hypothesised that incorporating additional evidence on molecular phenotypes could help to elucidate potentially novel loci which are likely to be identified as sample sizes of future GWAS increase. Although the observed effects of these loci on traits alone do not meet the conventional GWAS threshold, we took evidence of colocalization (again defined as a combined PPA >= 80%) at these loci as novel evidence implicating them in disease which can be used to prioritise them for future evaluation.

### Functional informatics

#### Pathway analysis

For all scenarios where GWAS trait and gene expression colocalize based on a combined PPA of >= 80%, we compiled a list of associated genes for each trait. Where multiple genes at a region provided a PPA >= 80% for the same GWAS SNP, we took forward the gene with the highest PPA. Pathway analysis was then undertaken with a gene list for each neurological trait using ConsensusPathDB^23^. This was to investigate whether multiple associated genes in our analysis reside along established biological pathways more than we would expect by chance.

#### Tissue-specific analysis

We also investigated whether any genes detected in our analysis were predominantly expressed in brain tissue using 3 RNA-seq datasets; the Human Protein Atlas (HPA)^24^, the Genotype-Tissue Expression project (GTEx)^25^ and the Mouse ENCODE project^26^. We used the ‘TissueEnrich’ R Package to identify evidence of enrichment based on 3 definitions^24^:

- **Tissue Enriched:** Genes with an expression level greater than 1 (TPM or FPKM) plus at least 5-fold higher expression levels in a particular tissue when compared to all other tissues.
- **Group Enriched:** Genes with an expression level greater than 1 (TPM or FPKM) plus at least 5-fold higher expression levels in a group of 2–7 tissues when compared to other tissues not considered to be ‘Tissue Enriched’.
- **Tissue Enhanced:** Genes with an expression level greater than 1 (TPM or FPKM) plus at least 5-fold higher expression levels in a particular tissue compared to the average levels in all other tissues, not considered to be either ‘Tissue Enriched’ or ‘Group Enriched’.

For each dataset we were only interested in genes predominantly expressed in brain tissue i.e. ‘Cerebral Cortex’ in HPA, ‘Brain’ in GTEx and ‘Cerebellum’,’Cortex’ or ‘E14.5 Brain’ in the Mouse ENCODE project. Heatmaps to illustrate enrichment across all possible tissues from these datasets were generated using the ‘ggplot’ R package’^27^.

#### Orienting directions of effect between molecular traits and regulatory region annotation

We oriented the direction of effect between transcriptional and epigenetic traits for detected associations; firstly between gene expression and DNA methylation and then between gene expression and histone acetylation. For associations with evidence of colocalization between the two traits being assessed, we evaluated whether the lead SNP was correlated with molecular traits in the same direction using coefficients from the xQTL resource. We applied the hypergeometric test to investigate whether there was an enrichment of a particular direction of effect between molecular traits more than we would expect by chance. Background expectations were calibrated using randomly selected lead SNPs across the genome that were associated with both proximal gene expression and DNA methylation (P < 1.0 × 10^−04^). Permutation testing was applied for 10,000 iterations by sampling the same number of SNPs being evaluated.

Lastly, we obtained regulatory data from the Roadmap Epigenetics Project^28^ from 10 different types of brain tissue. We used BEDtools^29^ to evaluate whether lead SNPs, CpG sites and histone peaks with evidence of colocalization from our study reside within promoters, enhancers and histone marks using these datasets. All analyses in this study were undertaken using R (version 3.31).

## Results

### Colocalization between gene expression, DNA methylation and histone acetylation with risk loci for 5 neurological traits and diseases

We applied the moloc method at loci with a trait-associated SNP (P < 5.0 × 10^−08^) using findings from 5 large-scale GWAS^15–18^ and molecular datasets (eQTL, mQTL and haQTL) derived from brain tissue^14^. Across the 5 neurological traits we identified a total of 66 colocalization associations with GWAS loci and gene expression (Supplementary Tables 2–6). Of these, 40 provided evidence of colocalization with an epigenetic trait also. Altogether, 4 genetic loci colocalized with a complex trait and all three of the molecular phenotypes (gene expression, DNA methylation and histone acetylation). Figure 1 illustrates these associations for neuroticism and insomnia, whereas plots for the remaining traits can be located in Supplementary Figure 1.

**Table 1.**
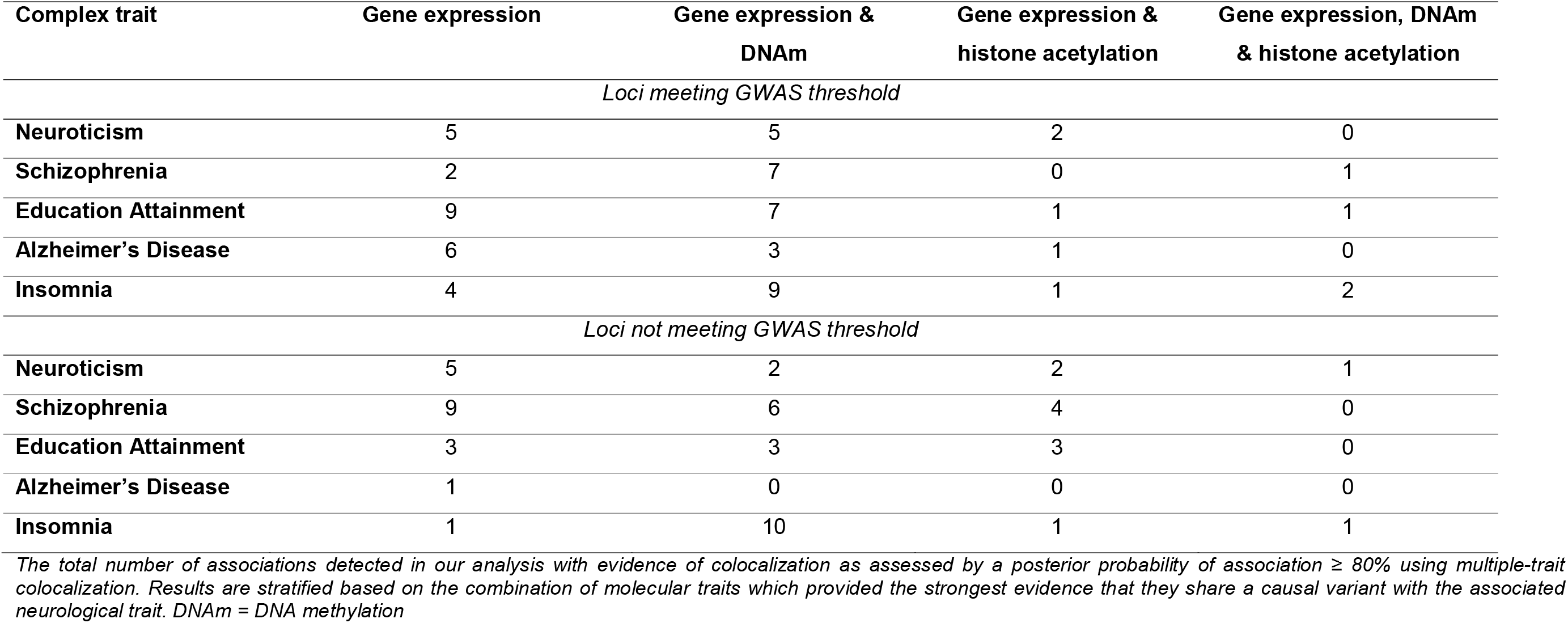
Number of associations with evidence of colocalization between molecular and neurological and molecular traits

| Complex trait | Gene expression | Gene expression & DNAm | Gene expression & histone acetylation | Gene expression, DNAm & histone acetylation |
| --- | --- | --- | --- | --- |
| *Loci meeting GWAS threshold* |  |  |  |  |
| Neuroticism | 5 | 5 | 2 | 0 |
| Schizophrenia | 2 | 7 | 0 | 1 |
| Education Attainment | 9 | 7 | 1 | 1 |
| Alzheimer's Disease | 6 | 3 | 1 | 0 |
| Insomnia | 4 | 9 | 1 | 2 |
| *Loci not meeting GWAS threshold* |  |  |  |  |
| Neuroticism | 5 | 2 | 2 | 1 |
| Schizophrenia | 9 | 6 | 4 | 0 |
| Education Attainment | 3 | 3 | 3 | 0 |
| Alzheimer's Disease | 1 | 0 | 0 | 0 |
| Insomnia | 1 | 10 | 1 | 1 |

*The total number of associations detected in our analysis with evidence of colocalization as assessed by a posterior probability of association $\geq 80\%$ using multiple-trait colocalization. Results are stratified based on the combination of molecular traits which provided the strongest evidence that they share a causal variant with the associated neurological trait. DNAm = DNA methylation*

**Figure 1.**
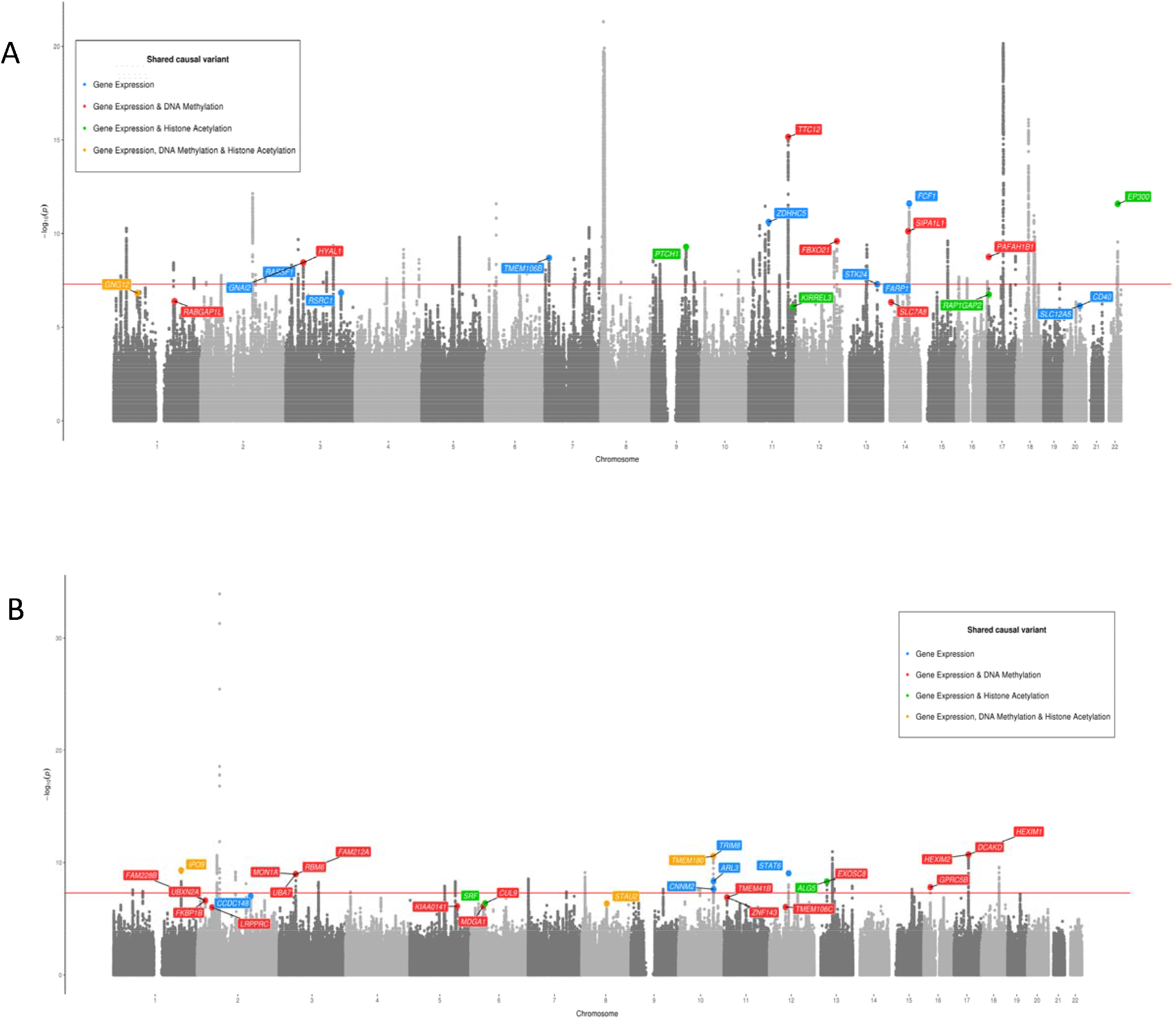
Manhattan plots for (A) Neuroticism and (B) Insomnia. Shared causal variants with traits are represented for the following scenarios; Gene expression (blue), Gene expression & DNA methylation (red), Gene expression & histone acetylation (green) and Gene expression, DNA methylation and histone acetylation (yellow). The genome-wide significance threshold (P < 5 × 10^−08^) is illustrated in red.

We identified evidence of colocalization between neurological and molecular traits at loci previously reported as well as novel findings. For example, we were able to replicate findings reporting that the expression of *KLC1* colocalises with schizophrenia risk and DNA methylation^13^ (combined PPA=97.9%). There were several other loci associated with schizophrenia that have been previously reported to colocalize with molecular traits (such as *CNNM2* and *PRMT7*^13^), as well as a several other genes where epigenetic mechanisms have not been previously detected to play a role in schizophrenia risk (such as *TSNARE1* and *ADOPT1*) (Supplementary Table 3).

There were also novel associations with molecular phenotypes amongst the other neurological traits. For instance, we uncovered evidence suggesting that neuroticism, gene expression and DNA methylation shared a CCV at the *PAFAH1B1* locus (combined PPA=89.9%). Figure 2A illustrates the overlapping distributions of effects on each of these traits for variants at this region. We also observed evidence of colocalization for several genes at the *APOE* locus that were associated with Alzheimer’s disease. This included *TOMM40*, where results suggested there was also evidence of colocalization with DNA methylation (combined PPA=99.3%). However, given the extensive linkage disequilibrium at this region, findings should be interpreted with caution^30^.

**Figure 2.**
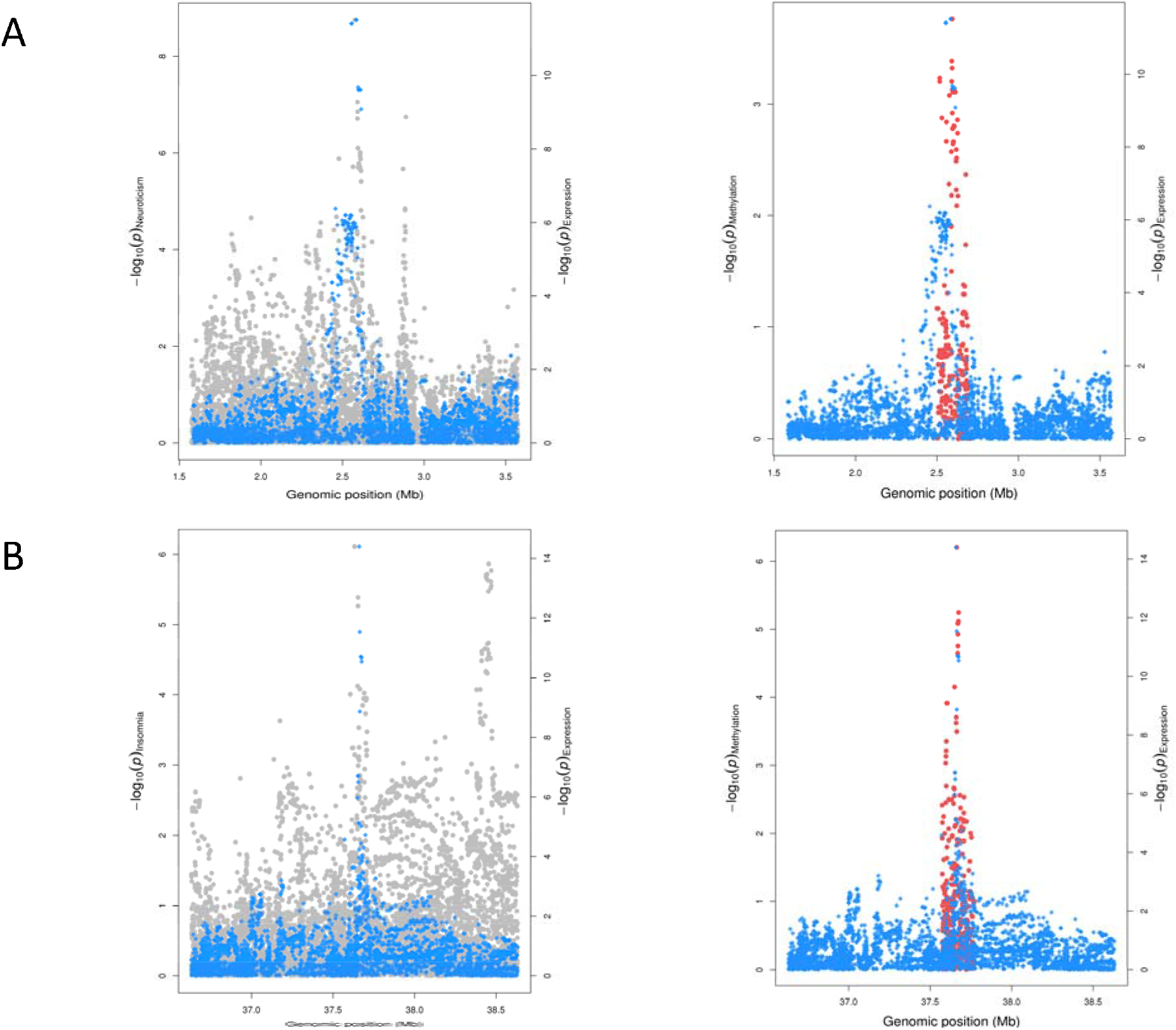
Regional association plots illustrating colocalizations for the *PAFAH1B1* gene (A) and *MDGA1* gene (B) with neuroticism and insomnia respectively. Effects for genetic variants on complex traits and gene expression were available within a 1Mb distance of the lead variant at each locus, whereas effects on DNA methylation levels were confined to a 5kb distance.

### Elucidating novel genes that may influence neurological traits

We also applied our analytical pipeline to uncover potentially novel loci using a less stringent threshold (P≤1.0×10^−06^). In this analysis we identified 52 loci where neurological traits and gene expression share a CCV, of which 33 provided evidence that these variants may also influence epigenetic traits (Figure 1; Supplementary Figure 1). Our incentive for undertaking this analysis was that GWAS analyses may not identify evidence of association using observed effects on neurological traits alone. However, by integrating evidence that SNPs at these loci also influence molecular traits derived from a relevant tissue type we aimed to uncover novel loci in disease/trait variation. Table 1 provides an overview of the number of associations detected in our analysis.

As an example of this, there was evidence that insomnia risk and molecular traits share a CCV at the *MDGA1* locus (combined PPA=85.8%). However, the p-value for the lead SNP at this region did not reach conventional GWAS thresholds (P = 7.7 × 10^−07^), suggesting that it would have been potentially overlooked based on GWAS evidence alone. As a validation of this finding, we found that a recent GWAS of insomnia with a larger sample size has found strong evidence of association at the *MDGA1* locus which survives conventional corrections (P=4.0×10^−12^)^31^. Figure 2B illustrates the overlapping distribution of effects for genetic variants at *MDGA1* on insomnia, gene expression and DNA methylation.

We identified several other instances from our analysis of loci with evidence of colocalization that have recently been detected by GWAS, suggesting that our analytical pipeline is valuable in terms of detecting novel findings. For example, we found that expression of the *CD40* and *SLC12A5* genes colocalize with risk of neuroticism. Both genes have subsequently been identified as associated with neuroticism at genome-wide significance in a GWAS meta-analysis^32^. Additionally, a recent large GWAS of educational attainment identified several genetic variants not previously found to reach genome-wide significance that we found to colocalize with molecular traits for the following genes: *DNAJB4, RERE, Corf73, DHX30, CD164* and *GLCC11^33^*.

### Pathway and tissue-specific enrichment analysis

Pathway analysis was conducted using ConsensusPathDB^23^ to investigate whether any sets of genes for each neurological trait and disease reside along the same biological pathway (Supplementary Table 12). Amongst findings there was evidence that genes associated with neuroticism in our analysis (*SLC12A5, GNAI2* and *GNG12*) reside on the GABAergic synapse pathway (enrichment P=2.34×10^−4^).

Our tissue-specific analysis indicated that various genes with evidence of colocalization are predominantly expressed in the brain. *SLC12A5, KLC1* and *KIRREL3* are expressed specifically in the cerebral cortex using data from the Human Protein Atlas^24^, whereas *MDGA1* was strongly expressed within brain tissue using data from the GTEx^25^ project. *RAP1GAP2* was predominantly expressed within cortex tissue using findings from the Mouse ENCODE project, amongst other loci are enriched in brain tissue based on this dataset (Supplementary Table 13, Supplementary Figure 2).

We observed enrichment of an inverse relationship between DNA methylation and gene expression across loci which provided evidence of colocalization for these molecular traits (P = 1.98 × 10^−03^), supporting previous evidence implicating DNA methylation as a transcriptional repressor^34^ (Supplementary Table 14). This effect appeared to be driven by CpG sites located near the transcription start site of genes, as 11 of the 13 sites located at these regions were inversely correlated with gene expression (84.6%). Performing the same analysis except with gene expression and histone acetylation suggested there was weak evidence of enrichment for a directional relationship (P=0.37). The regulatory annotations within brain tissue datasets for lead SNPs, CpG sites can be found in Supplementary Tables 15–17.

## Discussion

In this study we have conducted an integrative analysis of GWAS and molecular QTL data to uncover mechanistic insight into the biological pathways underlying complex neurological traits. We identified 118 colocalization associations between neurological traits and gene expression, with 73 of these associations additionally colocalizing with proximal DNA methylation and/or histone acetylation in brain tissue. Out of the 118 associations, 52 were potentially novel loci which did not meet genome-wide significance corrections but colocalized with molecular traits. Notably, several of these potentially novel loci have recently been validated by larger GWAS^31–33^, suggesting that other findings in our study are likely to be identified by GWAS as study sizes increase. Our findings should help future studies prioritise candidate genes and putative epigenetic mechanisms for functional follow-up analyses.

Applying our analysis pipeline to GWAS loci associated with neurological traits and disease (i.e. P<5×10^−08^) replicated previous findings reported by functional studies. For instance, our findings are consistent with an in-depth evaluation of the *KLC1* locus^3^. Variation at *KLC1* provided strong evidence of colocalization in our study (combined PPA=97.9%), where the highest individual posterior probability suggested both gene expression and DNA methylation may be involved along the causal pathway to schizophrenia risk. This result also supports findings from an epigenome-wide association study implicating DNA methylation as potentially playing a role in schizophrenia risk at this locus^35^. Furthermore, Hi-C interactions have been identified at the promoter region of *KLC1* within brain tissue which further helps validate the putative regulatory mechanism implicated by our results^36^.

Amongst other established GWAS loci, there was evidence suggesting that expression of the *TOMM40* gene and DNA methylation may play a role in Alzheimer’s disease. An exploratory analysis has found that regulatory element methylation levels in the *TOMM40-APOE-APOC2* gene region correlate with Alzheimer’s disease^37^. However, there is also evidence that, although SNPs at this region are known to influence Alzheimer’s disease, gene expression and DNA methylation, they may be attributed to different causal variants^38^. Moreover, there is a complex linkage disequilibrium structure at this region^30^, suggesting that further analysis is required to fully understand the mechanisms underlying this association.

We were also able to identify evidence of colocalization at GWAS loci that have not been linked previously by functional analyses or integrative studies harnessing molecular traits. For instance, the underlying biology explaining a GWAS association with neuroticism on chromosome 17 (lead SNP = rs12938775) has yet to be thoroughly evaluated. Our findings suggest that *PAFAH1B1* may be the likely causal gene at this locus, as well as implicating the involvement of DNA methylation along the causal pathway to neuroticism susceptibility as well (combined PPA=90.0%). *PAFAH1B1* (also known as *LIS1*) is involved in neuronal migration, the process by which different classes of neurons are brought together so that they can interact appropriately^39^. Functional evaluations of how changes in DNA methylation may influence neurological function at loci such as this may prove valuable in understanding epigenetic contributions to disease susceptibility. Moreover, doing so will help improve the accuracy of early disease prognosis.

As well as helping characterize associations detected by GWAS studies, we have also uncovered evidence for many novel genes which may influence neurological trait variation and therefore represent promising candidates for future endeavours. The association with insomnia risk at the *MDGA1* locus is an example of this, particularly given that it has recently been validated by a large-scale GWAS^31^. Furthermore, our results may provide functional insight into this association, by suggesting that *MDGA1* may be the responsible causal gene and that DNA methylation may also play a role in disease risk at this locus (combined PPA=85.8%). Similar to *PAFAH1B1, MDGA1* has also been report to play a role in neuronal migration^40^ and based on our tissue-specific analysis is predominantly expressed in brain tissue.

*SLC12A5*, associated with neuroticism in our analysis (combined PPA=99.0%), was amongst other promising candidates which has yet to be discovered by GWAS. This gene encodes the neuronal KCC2 channel which plays a crucial role in fast synaptic inhibition^41^. *SLC12A5* was also amongst the genes associated with neuroticism in our analysis that resides along the GABAergic synapse pathway (along with *GNAI2* and *GNG12*). A recent study has suggested that GABAegic neurons are causally associated with risk of bipolar disorder^42^, a condition previously linked with higher global measures of neuroticism^43^. The association between *KIRREL3* and neuroticism (combined PPA=92.2%) is another finding that has yet to be identified by GWAS which warrants in-depth functional evaluation. *KIRREL3* regulates target-specific synapse formation and has been previously linked with neurodevelopmental disorders^44^. Our tissue-specific analysis suggests that both *SLC12A5* and *KIRREL3* are predominantly expressed in brain tissue.

In cases where gene expression was found to colocalize with DNA methylation, we observed evidence of enrichment for an inverse relationship between these molecular phenotypes. Such inverse correlations support established biology that DNA methylation plays a role in silencing gene transcription^45^. However, recently there has been conflicting reports concerning whether DNA methylation on its own is sufficient to lead to transcriptional repression of promoters^34, 46^. Further analysis investigating the epigenetic mechanisms identified by our study should prove valuable in fully understanding the role of DNA methylation in gene regulation.

In terms of limitations of this study, we recognise that integration of GWAS results with QTL data is limited by the sample sizes used to derive summary statistics, which is particularly noteworthy for QTL data available in brain. Analyses such as ours are often limited in this manner, particularly when analysing brain-related phenotypes with sample sizes typically in the order of hundreds^47^. It may be the case that replication in blood can provide greater power due to the larger sample sizes available. It has been shown that top cis-eQTL and mQTL are highly correlated between blood and brain tissues^47^. Future work could take advantage of this correlation and the higher power in these blood datasets.

There is also evidence that the expression of certain genes is both highly tissue and disease-specific^42^. Recently, it has been shown that both tissue-specific and tissue-shared eQTL provide a substantial polygenic contribution to various complex traits^26^. Further investigation into the tissue-specificity of our results could be interesting since the ROSMAP/Brain xQTL^14^ dataset comes specifically from the dorsolateral prefrontal cortex region of the brain. Analysis of effects in other regions of the brain may be interesting to potentially identify disease relevant regions. We were also limited as the mQTL data was confined to 5kb windows affecting the coverage we could get within a genomic region. Whilst the nature of this mQTL dataset means we may have missed some true effects, it also means we are unlikely to have identified false positives. It is also worth noting that as the number of molecular studies increases, so too does the likelihood of detecting incidental QTLGWAS overlaps^3^. Hence, developments concerning robust methods in colocalization should prove to be extremely valuable and important for future research.

By integrating GWAS findings with data concerning brain cortex-derived molecular phenotypes, we have helped uncover putative epigenetic and transcriptomic drivers of neurological function and disease. Our work has focused on the prioritisation of GWAS hits and uncovering potentially novel loci which are likely to influence various complex neurological traits. The analytical framework applied in our study can be harnessed to help to characterise biological mechanisms for a wide variety of different traits and disease.

## Funding

This study and C.H were supported by a 4-year studentship fund from the Wellcome Trust Molecular, Genetic and Lifecourse Epidemiology Ph.D. programme at the University of Bristol (108902/B/15/Z). The MRC Integrative Epidemiology Unit receives funding from the UK Medical Research Council and the University of Bristol (MC_UU_00011/4 and MC_UU_00011/5). T.G.R is a UKRI Innovation Research Fellow (MR/S003886/1). The authors declare no conflicts of interest.

## Acknowledgements

We are grateful to the developers of the Brain xQTL resource (http://mostafavilab.stat.ubc.ca/xqtl/) and authors of the GWAS studies cited in this study for making their summary statistics publicly available. We would also like to thank Claudia Giambartolomei, the lead developer of ‘moloc’, for all her time and help with questions regarding our analysis plan. This publication is the work of the authors and T.G.R. will serve as guarantor for the contents of this paper.

